# Alteration of functional connectivity in the cortex and major brain networks of non-human primates following focused ultrasound exposure

**DOI:** 10.1101/2023.02.16.528741

**Authors:** D Liu, F Munoz, S Sanatkhani, A N Pouliopoulos, E Konofagou, J Grinband, VP Ferrera

## Abstract

Focused ultrasound (FUS) is a non-invasive neuromodulation technology that is being investigated for potential treatment of neurological and psychiatric disorders. Focused ultrasound combined with microbubbles can temporarily open the intact blood-brain barrier (BBB) of animals and humans, and facilitate drug delivery. FUS exposure, either with or without microbubbles, has been demonstrated to alter the behavior of non-human primates, and previous work has demonstrated transient and long-term effects of FUS neuromodulation on functional connectivity using resting state functional MRI. However, it is unknown whether opening the BBB affects functional connectivity differently than FUS alone. Thus we applied FUS alone (neuromodulation) and FUS with microbubbles (BBB opening) in the dorsal striatum of lightly anesthetized non-human primates, and compared changes in functional connectivity in major brain networks. We found different alteration patterns between FUS neuromodulation and FUS-mediated BBB opening in several cortical areas, and we also found that applying FUS to a deep brain structure can alter functional connectivity in the default mode network and frontotemporal network.

## Introduction

Focused ultrasound (FUS) is a non-invasive brain stimulation technology that is in early stages of translation for clinical applications in psychiatry and neurology. While high intensity focused ultrasound is capable of producing lesions (Ter Haar & Coussios, 2007), low intensity burst mode FUS can deliver energy non-invasively into specific deep brain regions without trauma (Kubanek, 2018; Munoz et al. 2022). These properties make it an attractive alternative to other neuromodulation technologies, such as transcranial direct current stimulation (tDCS) (Nitsche et al., 2008) and transcranial magnetic stimulation (TMS) (Walsh & Cowey, 2000), both of which primarily target superficial brain regions. Animal studies have shown that FUS could be used for noninvasive cortical and subcortical brain stimulation with sub-millimeter focus (Tufail et al., 2010), by inducing excitatory or inhibitory effects in the central or peripheral nervous system, depending on the pulsing regime (Blackmore et al., 2019). FUS can also alter behavior in non-human primates (NHPs) (Deffieux et al., 2013; Fouragnan et al 2019; Munoz et al., 2022; Banaie Boroujeni et al., 2022).

The mechanism by which FUS can affect the functional connectivity of brain regions and networks has been studied using resting state fMRI, a method based on the blood oxygenation level dependent (BOLD) contrast. Sanguinetti et al (Sanguinetti et al., 2020) found application of FUS to the prefrontal cortex can alter the functional connectivity and improve mood in humans. In NHPs, recent studies used resting state fMRI to evaluate long-lasting or “offline” effects of FUS exposure on subcortical or deep cortical regions, including amygdala, anterior cingulate cortex (Folloni et al., 2019), medial frontal cortex (Bongioanni et al., 2021), and supplemental motor cortex (Verhagen et al., 2019). Munoz et al (Munoz et al., 2022) found that 2 minutes of FUS applied to the dorsal striatum can alter patterns of functional connectivity in the NHP brain that can last for hours and can be correlated with improved cognition.

FUS combined with microbubbles has been used to reversibly permeabilize the blood-brain barrier (BBB) and facilitate drug delivery both in animal and human studies (McDannold et al., 2012; Meng et al., 2019a), and recent work has demonstrated that FUS exposure with microbubbles in the striatum can improve response speed and accuracy during visual-motor tasks in NHPs (Chu et al., 2015; Downs et al., 2017). Using resting state fMRI, Todd et al (Todd et al., 2018) found that FUS-induced BBB opening can disrupt the functional connectivity between inter-hemispheric regions in rats. Meng et al (Meng et al., 2019b) found transient functional connectivity reductions with FUS BBB opening in the frontal lobe of patients with Parkinson’s disease.

Although both FUS and FUS with microbubbles (BBB opening) have been shown to modulate neural activity, the impact of FUS-induced BBB opening on functional connectivity is not well understood. Changing the permeability of the BBB may alter blood perfusion, nutrient absorption, or tissue oxygenation near the BBB opening and may affect neural function differently than direct neuromodulation. No prior studies have investigated the effects of FUS-mediated BBB opening on functional connectivity in non-human primates. Investigating how BBB opening differs from direct neuromodulation is needed to understand how BBB opening affects cognitive performance and to establish a baseline for evaluating the outcomes of FUS-mediated drug delivery in the treatment of neurological and psychiatric disorders. The objective of this study was to compare the effects of applying FUS (neuromodulation) and FUS combined with microbubbles (BBB opening) to the NHP dorsal striatum in order to determine how changes in resting state functional connectivity of major brain networks differs between the two approaches.

## Results

### Numerical simulation and post-FUS exposure confirmation

For precise targeting and energy deposition of FUS during the experiments, acoustic and biothermal simulations of a single element transducer (ROC=64 mm) were conducted using Matlab k-Wave toolbox and NHP skull CT (computer tomography) model. For burst mode ultrasound protocol described in Fig. 1a (500 kHz central frequency, 10 ms pulse duration, 2 Hz pulse repetition frequency, 2% duty cycle and 2 min total sonication), both the peak negative pressure (PNP) and temperature distribution are shown in Fig. 1b. The PNP of the target at the right caudate can reach above 800 kPa, while maximum temperature in the target is within 37.5°C. More ultrasound energy was deposited in the skull due to its high absorption coefficients compared to soft brain tissue, and the temperature elevation reached 3-4°C above baseline body temperature. Fig. S1 demonstrates that the maximum temperature of the skull, muscle, and brain were 41.2°C, 39.7°C and 38.2°C respectively, within the safety limits of the current guideline (Moyano et al., 2022).

**Fig. 1.**
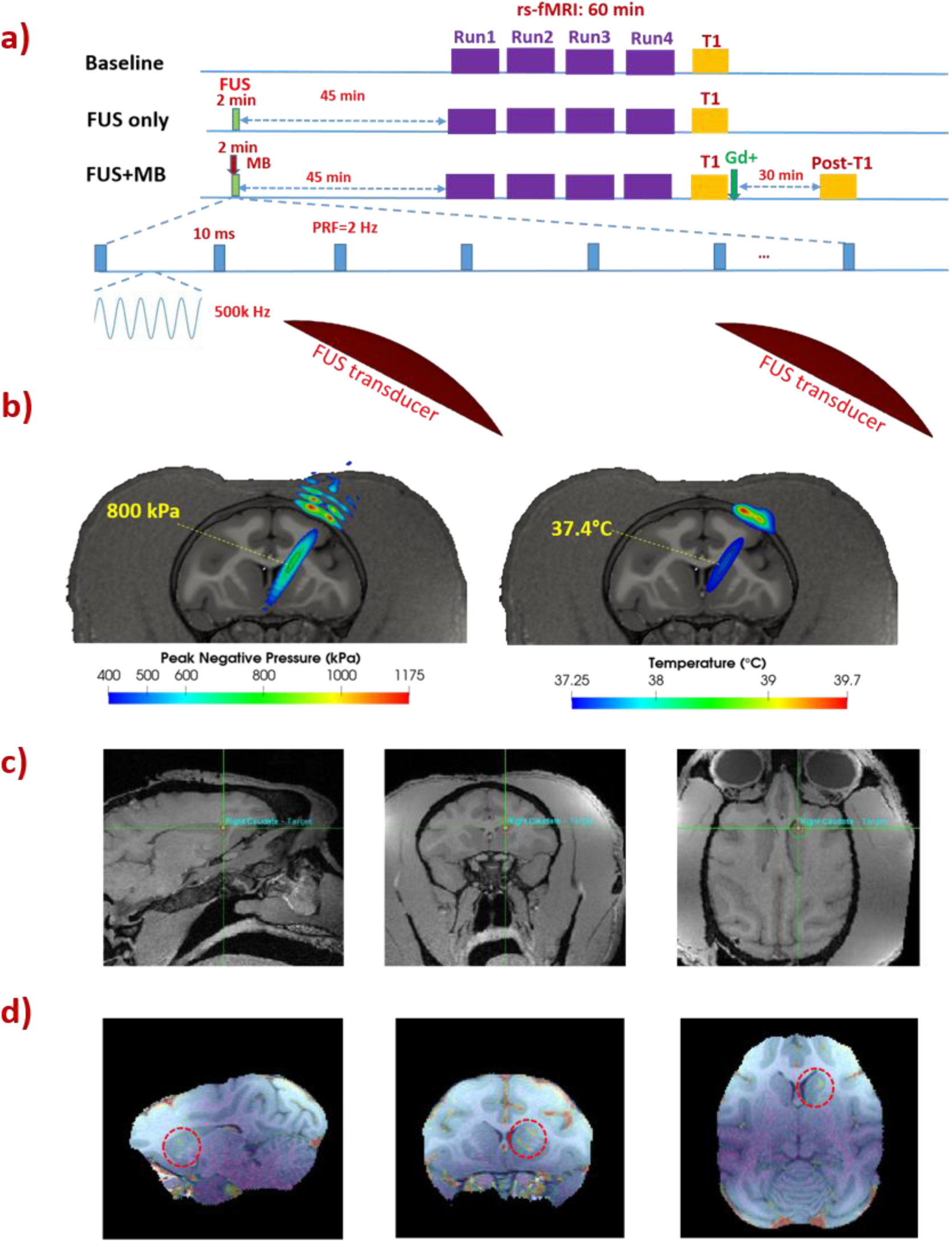
MRI and FUS experimental setup, pre-treatment planning and post-treatment confirmation: a) MRI and FUS scheme including baseline, FUS exposure (FUS only) and FUS with microbubbles (FUS+MB); b) FUS Acoustic (derated PNP ∼800 kPa in the target) and biothermal simulation (∼0.5°C elevation in the target); c) FUS targeting on right caudate using the navigation device and d) Post Gadolinium T1 images as a confirmation for success of FUS-BBB opening, with BBB opening regions shown in red circle.

The planning and targeting of FUS exposure with and without microbubbles was achieved using the combined FUS and Brainsight neuro-navigation system. As shown in Fig. 1c, the FUS energy was delivered to target the region located in the caudate nucleus in the right hemisphere. For NHPs undergoing FUS exposure with microbubbles, Gadolinium enhanced structural scans were acquired, with hyper-intense regions indicating the sites of FUS-BBB opening shown in Fig. 1d. A good match between the planned target (Fig. 1c) and the actual BBB opening site (Fig. 1d) was found.

### Effects on the striatum and cortical regions

To quantify the functional connectivity due to FUS neuromodulation and BBB opening, a region of interest (ROI) with 3.5 × 3.5 × 3.5 mm^3^ voxel at the right caudate was first chosen as a seed. The seed correlation maps of baseline, FUS, and FUS with microbubbles, as well as the difference maps are shown in Fig. 2a, overlaid on the standard D99 NHP template (Reveley et al., 2017). With FUS exposure in the right caudate, the functional connectivity is found between caudate and insular cortex (IC: regions of Ial, Iapl and Id in both hemispheres shown in Fig. 2b), and between caudate and temporal cortex (areas RT, RTp and TGdd in both hemispheres shown in Fig. 2b) while the functional connectivity is reduced between caudate and motor cortex (MC: areas F1, F2, F3 and F4 in both hemispheres shown in Fig. 2b), and between caudate and somatosensory cortex (SSC: areas 1, 2, 3a and 3b in both hemispheres shown in Fig. 2b). In contrast, when applying FUS with microbubbles, the alteration of functional connectivity does not show the similar pattern as above. Instead, a stronger activation of functional connectivity is found between caudate and medial prefrontal cortex (mPFC: areas 8Bm, 9m, 9d and 10mr in both hemispheres shown in Fig. 2b). Statistical analysis on representative regions (area 9m of mPFC; area Ial of IC and area F1 of MC) were performed, with results of both Kruskal-Wallis test and post-hoc permutation test are shown in Fig. 2c. With FUS exposure in the right caudate, the activation of insular cortex (p<0.001 in right hemisphere and p<0.01 in left hemisphere, compared to baseline and FUS-BBB opening) and inhibition of the motor cortex (p<0.001 in both hemispheres, compared to other two conditions) is statistically significant, while with FUS-mediated BBB opening, significant changes of functional connectivity is only found within mPFC (p<0.01 compared to baseline).

**Fig. 2.**
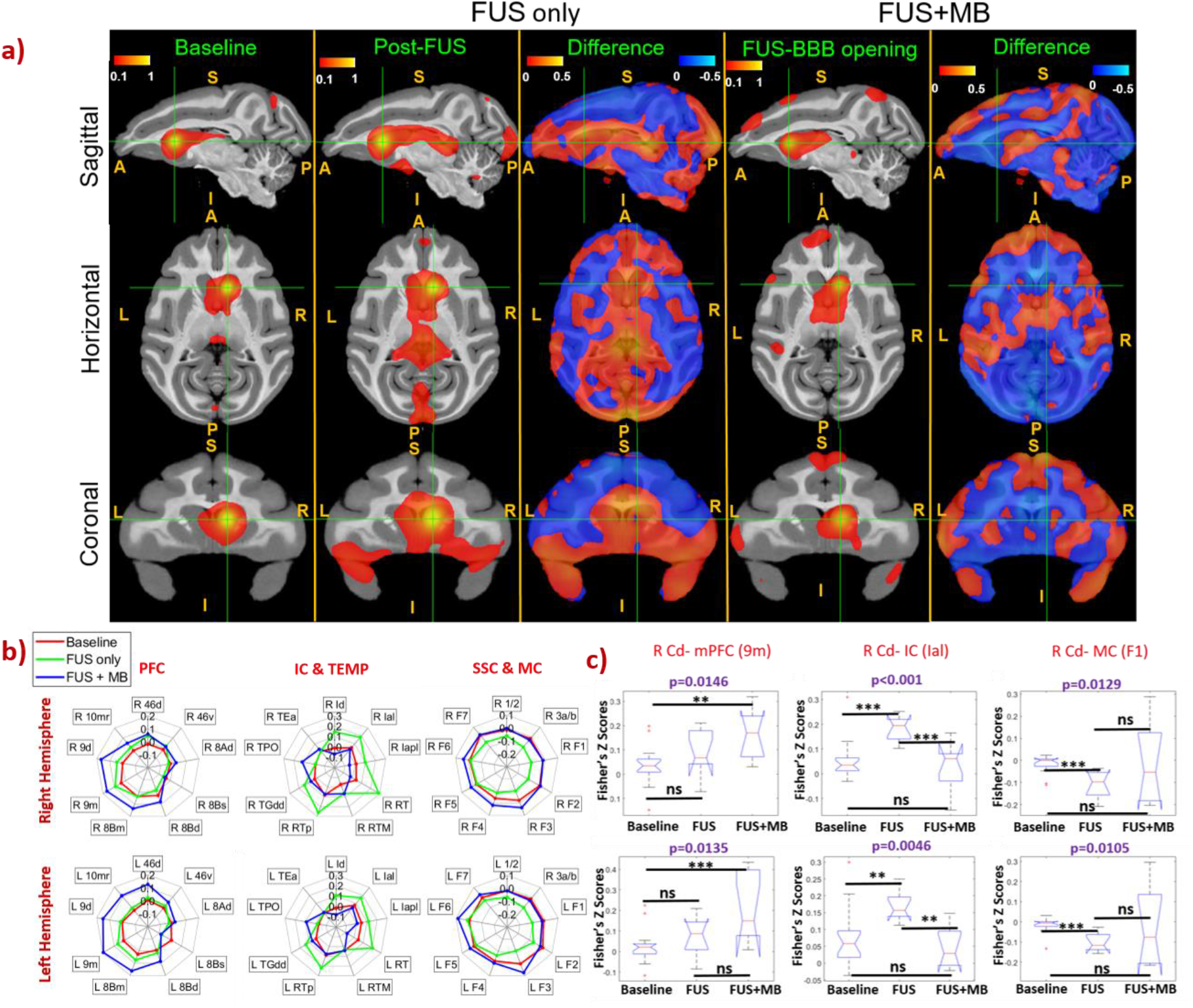
Seed-based connectivity and difference maps (a), radar charts (b) and statistical box plots of resting state fMRI without FUS (baseline) and after FUS (FUS only) and FUS with microbubbles (FUS+MB), indicating impact of FUS neuromodulation and FUS-mediated BBB opening on brain functional connectivity. Connectivity changes were found in the post-FUS and post-FUS BBB opening compared to the baseline. The red-yellow and blue-light blue color maps indicate Fisher z-score maps with seeds in the right caudate (FUS target).. Statistical analysis was performed using Kruskal-Wallis test, with p<0.05 representing a significant difference. Post-hoc permutation test (5000 resamples) was performed when there is a significant difference among the three conditions, and * denotes p<0.05, ** denotes p<0.01, *** denotes p<0.001 and ns denotes p>0.05. L: left, R: right, A: anterior, P: posterior, S: superior, I: inferior. PFC: prefrontal cortex, IC: insular cortex, TEMP: temporal cortex, SSC: somatosensory cortex, MC: motor cortex, R Cd: right caudate.

### Effects on the default mode network

To quantify the effect of FUS on the default mode network, an ROI of 3.5 × 3.5 × 3.5 mm^3^ voxel was placed in the area 8Ad of dlPFC within the default mode network, with the central coordinates of the ROI listing in Table S1. The average correlation maps of baseline, FUS and FUS BBB opening, as well as the difference maps were calculated with the results shown in Fig. 3a. Both FUS exposure and FUS exposure with microbubbles demonstrate an activation of the default mode network (Fig. 3a and Fig. 3b), a network connecting dorsolateral prefrontal cortex (dlPFC), posterior cingulate cortex (PCC) and posterior parietal cortex (PPC) of NHPs. Statistical analysis on representative nodes (area 8Ad in dlPFC, areas 31 and 23b in PCC and areas LIPd in PPC, area TPO in temporal lobe (TEMP)) was performed, with the results of Kruskal-Wallis test for comparison among the three conditions, and post-hoc permutation test for comparison between each two conditions, as shown in Fig. 4a. With either FUS or FUS combined with microbubbles, significant functional connectivity changes were found between dlPFC and the areas of PPC, PPC and TEMP. When placing the seed at area 23b of PCC, the connectivity between PCC and dlPFC was also found to be significantly enhanced (Fig. 4c). However, the functional connectivity due to FUS and FUS with microbubbles was not significantly different (Permutation test; p>0.05, shown in Fig. 4a).

**Fig. 3.**
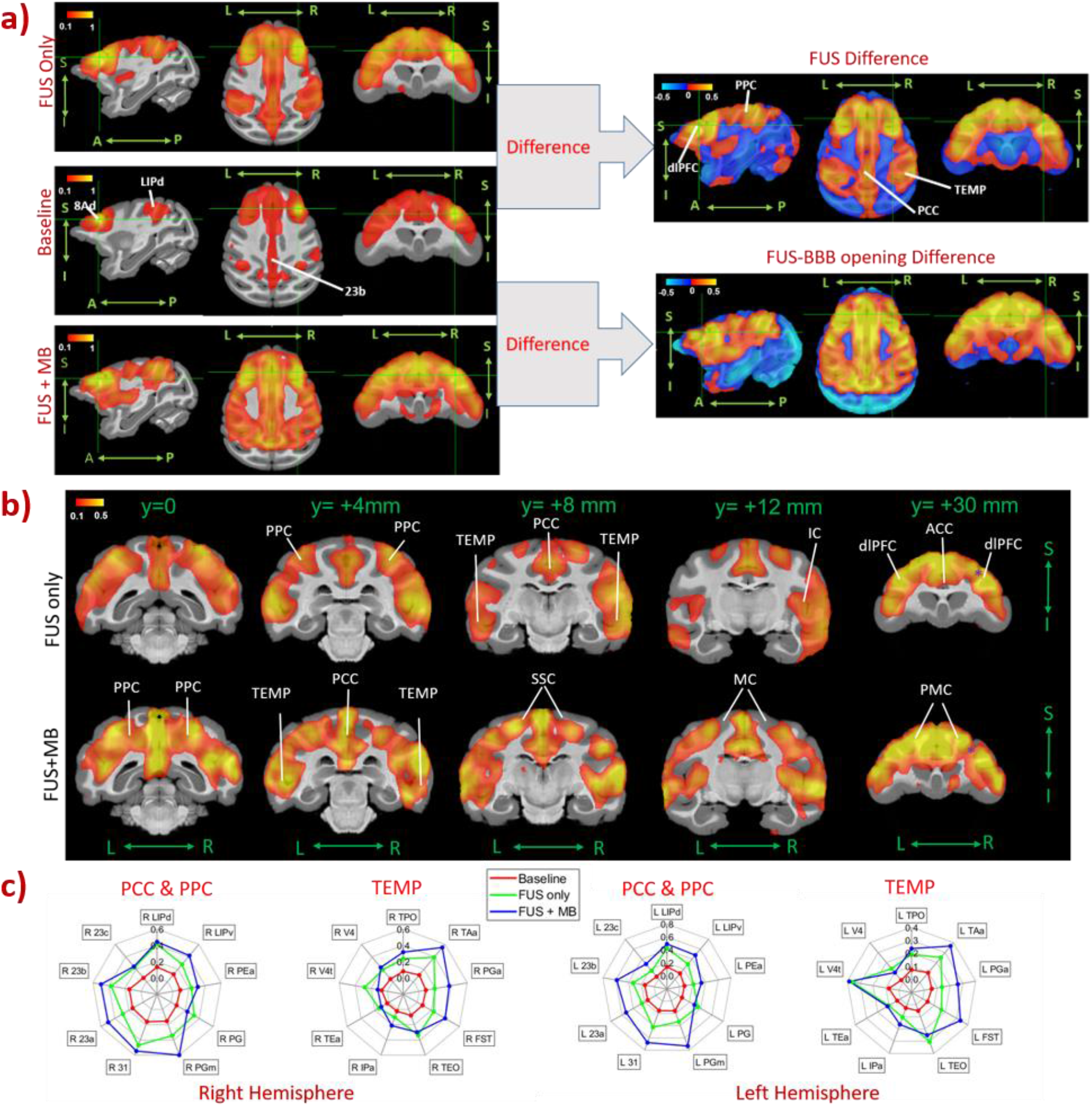
Seed-based connectivity maps (a), difference maps in coronal planes (b) and radar charts of resting state fMRI without FUS (baseline) and post-FUS (FUS only) and post-FUS with microbubbles (FUS+MB), indicating impact of FUS and FUS-induced BBB opening on brain functional connectivity. With the seed selected in 8Ad of dlPFC (green crossed point in (a)), Default mode network nodes were found significantly activated after FUS and FUS-BBB opening compared to the baseline. The red-yellow and blue-light blue color maps indicate Fisher z-score maps with seeds in right 46d. L: left, R: right, A: anterior, P: posterior, S: superior, I: inferior. ACC: anterior cingulate cortex, PCC: posterior cingulate cortex, PPC: posterior parietal cortex, TEMP: temporal cortex, IC: insular cortex, dlPFC: dorsal lateral prefrontal cortex, SSC: somatosensory cortex, PMC: primary motor cortex, MC: motor cortex.

**Fig. 4.**
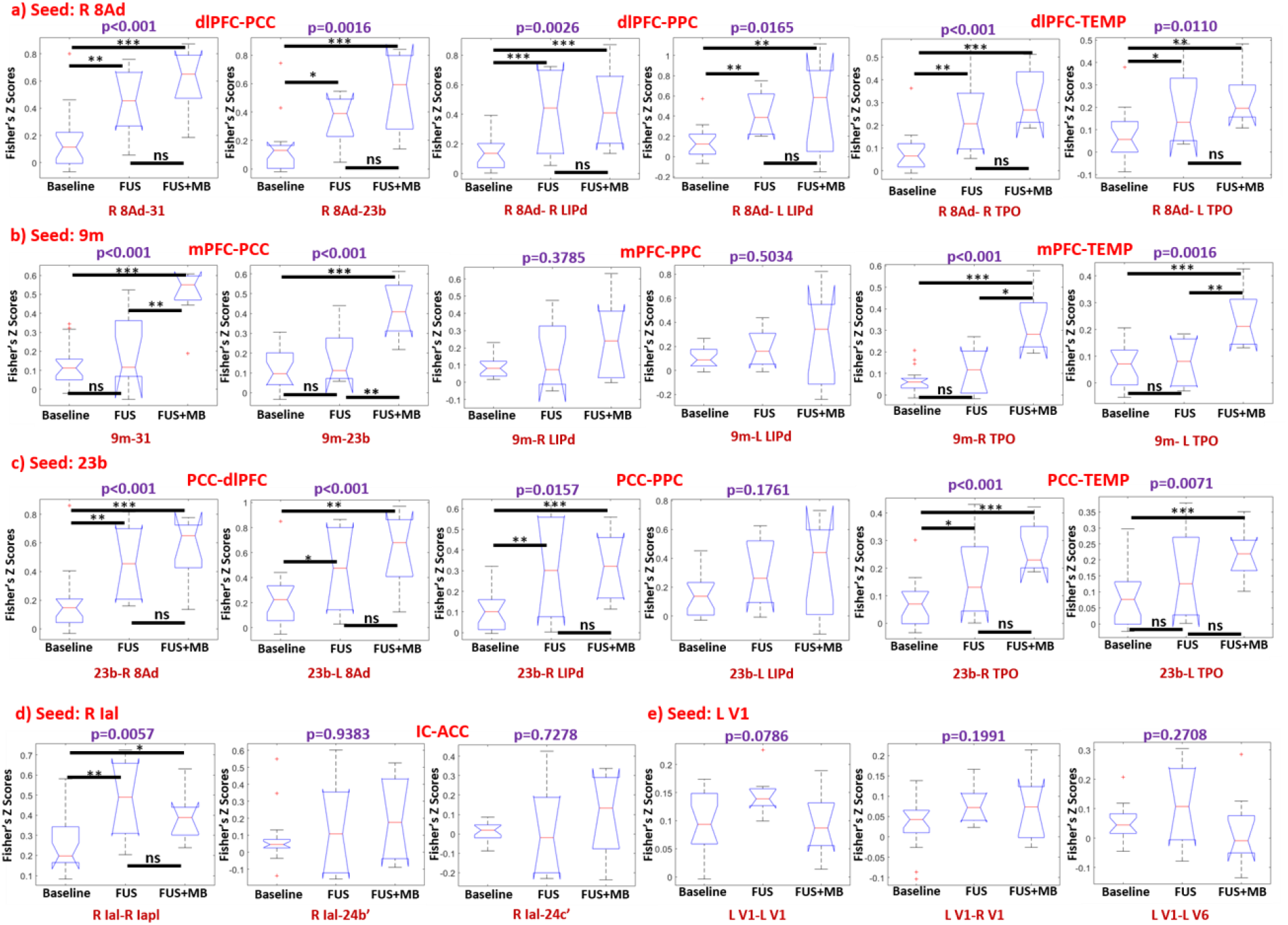
Seed-based correlation box plots with the seed ROI selected as right 8Ad (a), 9m (b), 23b(c), right Ial (d) and left V1 (e), indicating the changes functional connectivity among dlPFC, PCC, PPC, TEMP, IC and ACC with the conditions of baseline, FUS and FUS with microbubbles (FUS+MB). Statistical analysis was performed using Kruskal-wallis test, with p<0.05 representing a significant difference. Post-hoc permutation test (5000 resamples) was performed when there was a significant difference among the three conditions, and * denotes p<0.05, ** denotes p<0.01, *** denotes p<0.001 and ns denotes p>0.05. dlPFC: dorsal lateral prefrontal cortex, mPFC: medial prefrontal cortex, PCC: posterior cingulate cortex, PPC: posterior parietal cortex, TEMP: temporal cortex, ACC: anterior cingulate cortex, R: right, L: left

### Effects on the frontotemporal network and other networks

The effects of FUS neuromodulation and BBB opening on the frontotemporal network of the NHP, which connects the areas of mPFC and the temporal lobe, were evaluated. When selecting area 9m as a seed ROI in mPFC, only FUS combined with microbubbles demonstrates significant activation of functional connectivity between mPFC and regions of TEMP and PCC, as shown in Fig. S2 and Fig. 4b, while FUS (without microbubbles) failed to alter the functional connectivity significantly between mPFC and TEMP, compared to the baseline (FUS vs Baseline via post-hoc permutation test: p>0.05, shown in Fig. 4b).

While significant alteration of functional connectivity within DMN and FTN is found, the effect of FUS on other brain networks is not conclusive. For example, with the seed ROI of Ial, the average correlation between insular cortex and anterior cingulate cortex (ACC) with FUS-induced BBB opening is slightly higher than the correlations with the other two conditions, however, no statistically significant difference is found between FUS and FUS with microbubbles (Permutation test, p>0.05, shown in Fig. 4d). With the seed ROI chosen as primary visual cortex, no clear alteration of the functional connectivity is found between visual cortex and the other brain regions (Fig. S3), and no statistical difference is found among baseline, FUS and FUS with microbubbles (Kruskal-Wallis test, p>0.05, shown in Fig. 4e).

## Discussion

By comparing the “offline” resting state functional connectivity after applying 2 min FUS exposure with and without microbubbles in dorsal striatum of nonhuman primates, we found that FUS neuromodulation and FUS BBB opening affect patterns of functional connectivity differently. FUS neuromodulation can increase functional connectivity between caudate and insular cortex and decrease functional connectivity between caudate and motor cortex, while FUS-mediated BBB opening can increase functional connectivity between caudate and medial prefrontal cortex and the nodes within the frontotemporal network (Fig. 5b). The findings provide further evidence that FUS can be used as a neuromodulation technology to selectively modulate cortical and subcortical brain regions, which are commensurate with other studies (Folloni et al., 2019; Fouragnan et al., 2019; Verhagen et al., 2019). Specifically, significant alteration of functional connectivity between caudate and cortical regions were achieved in this study, using relatively lower average ultrasound intensity and slightly longer sonication compared to previous studies.

**Fig. 5.**
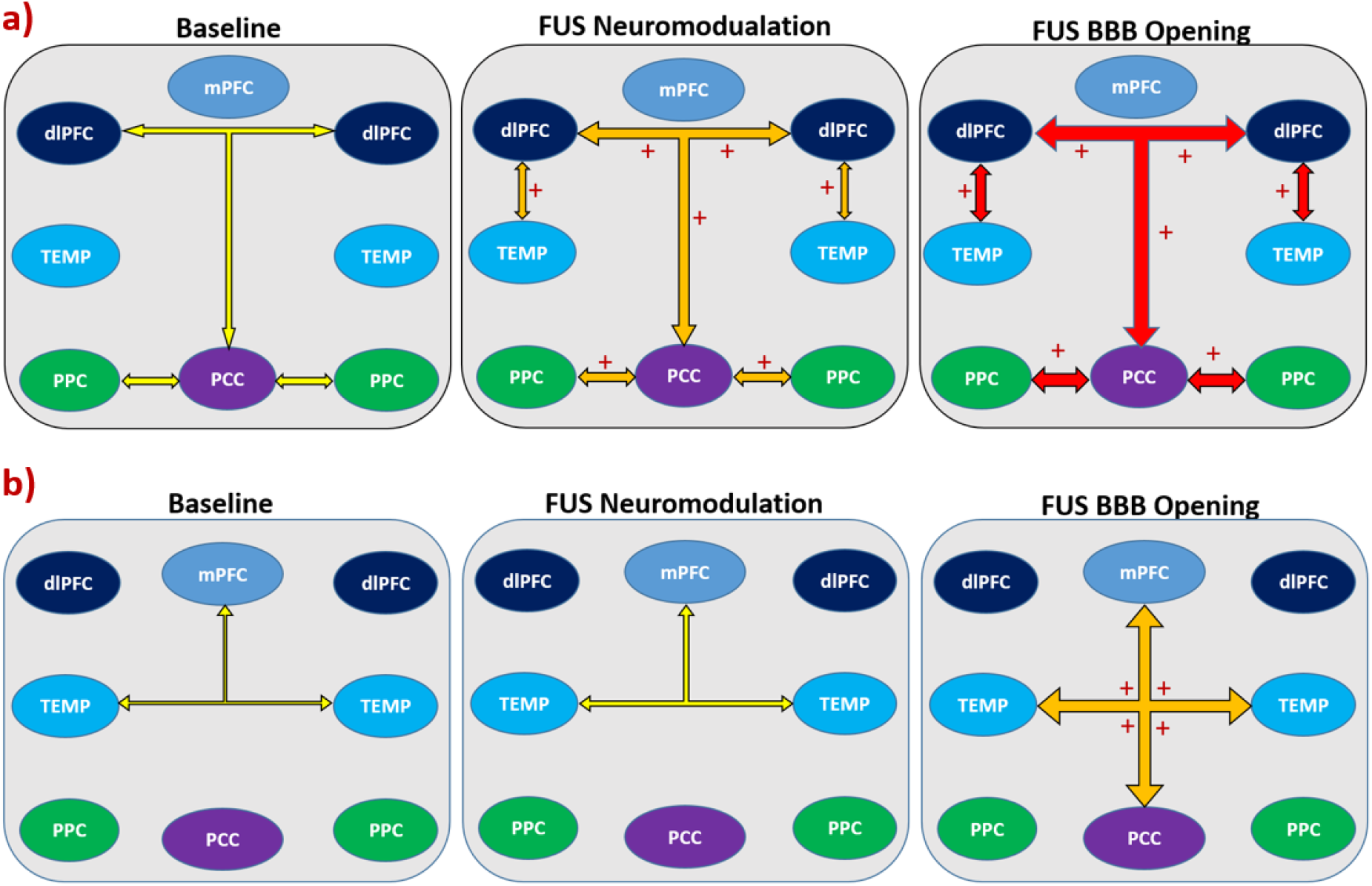
Summary of changes of functional connectivity on major brain networks among baseline, FUS neuromodulation and FUS-BBB opening: an activation of default mode network by both FUS neuromodulation and FUS-BBB opening and an activation of frontoparietal network by FUS-BBB opening. + indicates an activation with significant statistical difference. dlPFC: dorsal lateral prefrontal cortex, mPFC: medial prefrontal cortex, PCC: posterior cingulate cortex, PPC: posterior parietal cortex, TEMP: temporal cortex

Similar to recent findings in rodents (Todd et al., 2018) and humans (Meng et al., 2019b), the current study found that FUS-mediated BBB opening can temporarily alter functional connectivity in NHP. Currently, FUS-mediated BBB opening for drug delivery has demonstrated promising results in clinical trials, our study suggests that an accompanying neuromodulation effect may occur during the procedure of FUS-mediated BBB opening. Different patterns of functional connectivity may suggest the direct FUS and FUS BBB opening have different neuromodulation effects. This finding also reflects the results in visual-motor tasks in NHPs after applying FUS exposure with and without microbubbles: NHP improved accuracy along with a shorter response time after applying FUS exposure combined with microbubbles (Downs et al., 2017; Pouliopoulos et al., 2021), suggesting increased decision efficiency, while NHP showed more accuracy but with a longer response time after applying FUS exposure only, consistent with a speed-accuracy trade-off (Munoz et al., 2022). One possible explanation of the difference between FUS neuromodulation and FUS-induced BBB opening is that when BBB is opened by FUS combined with microbubbles in the target region, the blood oxygenation in that region may change as well (Todd et al., 2019). This may underlie changes in BOLD activation for resting state functional MRI. Future studies plan to address quantitative measurement of the blood oxygenation of FUS and FUS BBB opening using advanced MRI.

The study also demonstrated that applying FUS exposure in the caudate nucleus could activate the default mode network on lightly anesthetized NHP (Fig. 5a). Similar trends were found between FUS and FUS BBB opening, however we could not significantly differentiate the magnitude of functional connectivity of the two due to limited NHP experiments in this study. Findings from both FUS neuromodulation and FUS-mediated BBB opening may improve our understanding of the role of striatum in regulating cortical circuits, as well as the functional connection between striatum and default mode network. While the mechanism for external FUS stimulation in modulating the DMN via striatum is still unknown, one possible explanation may suggest striatum may “communicate” with cortical regions within DMN in a similar way with thalamus, which can both drive and modulate cortical regions (Sherman, 2016). A recent study in humans found that a reward task can enhance the connection between existing DMN and ventral striatum (Dobryakova & Smith, 2022), which also support that the striatum may play a role in regulating the DMN.

This study has several limitations. Firstly, awake and anesthetized NHP may have different patterns of functional connectivity (Xu et al., 2019), and isoflurane may affect the results of functional connectivity (Zhang et al., 2019), although in this study the same level (∼1.0 %) isoflurane was applied for all NHPs during fMRI collection to avoid the variation of BOLD activity due to varying anesthesia status. Functional connectivity was also reported to be related with age (Rao et al., 2021), and in this study, we did not rule out the effect of age on the results due to limited NHP resources. Another aspect that may affect variation of functional connectivity is the seed ROI we selected in this study (Lv et al., 2018). For example, when evaluating the effect of FUS on DMN, we selected some representative ROI’s in DMN following recent work evaluating DMN among primates (Mantini et al., 2011). Since the organization of DMN varies among non-human primates and humans (Garin et al., 2022), we tested different seeds (i.e. areas 8Ad, 46d, 8Bs in dlPFC, shown in Fig. S4) in this study to make sure the results are valid and robust.

In conclusion, this study compared the resting state functional connectivity on NHP after FUS neuromodulation and FUS-mediated BBB opening and found different alteration patterns in various cortical regions. Applying FUS to deep brain structures can alter functional connectivity in major brain networks such as the default mode network and frontotemporal network.

## Materials and methods

### Animal Preparation

A total of six adult male NHPs (N, P, O, Q, M and T, 6.9-12.3 kg) were used in the experiments. Two NHPs (P and O) were selected to apply FUS exposure and two NHPs (M and T) were selected to apply FUS exposure with microbubbles. All the six NHPs were scanned to acquire structural and functional MRI images. Prior to the ultrasound and MRI experiments, all the NHPs were firstly sedated with ketamine (10 mg/kg) and dexmedetomidine (0.02 mg/kg) and then were anesthetized with 1-1.5% isoflurane. The head skin of each NHP was shaved and the conductive gel was applied to achieve optimal ultrasound sonication. During the MRI experiments, the Iradimed 3880 MRI compatible monitoring system (Winter Springs, FL, USA) was used to wirelessly monitor the vital signs of NHPs including body temperature, electrocardiogram, oxygen saturation and respiratory CO2. All the NHP procedures were approved by the Institutional Animal Care and Use Committee (IACUC) of Columbia University.

### FUS neuromodulation and BBB opening

A single element FUS transducer (H-107, 500kHz frequency, 63.2 mm radius of curvature (ROC), 64 mm outer diameter (OD), Sonic Concepts, Bothell, WA) was driven by a functional generator (Aglient 33220A, Agilent Technologies, Santa Clara, CA) connecting to a 57dB radiofrequency power amplifier (500S06, E&I, Rochester, NY). The transducer cone was filled with degassed water using a water degassing system WDS105+ (Sonic Concepts, Bothell, WA), and was inflated to attach the head skin for efficiently delivering ultrasound energy to the target.

A FUS protocol described in previous studies (Karakatsani et al., 2017; Munoz et al., 2022) was adopted in order to achieve safe and efficient neuromodulation and BBB opening on NHPs, with detailed sonication parameters shown in Fig.1a. For the group with FUS neuromodulation, FUS generated by the single element FUS transducer was applied on NHPs O and P for 2 min with derated peak negative pressure (PNP) of 800 kPa, 2 Hz pulse repetition frequency (PRF), 10 ms pulse duration (PD) and 2% duty cycle. For the group with FUS-BBB opening, same FUS parameters but with derated PNP of 400 kPa were applied on NHPs M and T, and in-house manufactured microbubbles (MB) (4-5 *μm* diameter, 2.5× 10^8^ bubbles/kg) was injected intravenously through the saphenous vein of NHPs 10s after starting of the ultrasound sonication. The spatial peak temporal average intensities (Ispta) of the FUS neuromodulation and FUS-BBB opening are 156.9 mW/cm^2^ and 39.2 mW/cm^2^ respectively. In this way, a lower ultrasound intensity was used to eliminate the effects of neuromodulation but efficiently and safely enough to open the intact BBB of NHP, as reported in the previous literature (Pouliopoulos et al., 2021).

### Targeting and Numerical simulation

To achieve precise targeting of each NHP, a real time neuro-navigation system (BrainSight Vet System, Rogue Research Inc. Canada) in conjunction with the ultrasound system described in previous works was used (Wu et al., 2018). After calibrating the FUS transducer in water at room temperature, we set up a numerical model incorporating a FUS transducer (ROC=64 mm) adopted in this study and an NHP CT model, and both 3D acoustic and biothermal simulations were performed in Matlab k-wave toolbox (Treeby and Cox., 2010). For all the tissues including skull, brain and muscle, speed of sound and the density were set up following a linear relation with CT Hounsfeld units (Deffieux & Konofagou, 2010), frequency dependent attenuation coefficients were adopted based on previous measurement (Goss et al., 1979; Duck, 2013). The detailed acoustic and thermal parameters for numerical simulation were listed in Table S2 (Hasgall et al., 2022). The initial temperature of the tissue was set as 37°C, and the Dirichlet boundary condition was applied. To prevent the skull from heating, the cone water temperature was set as 22°C to provide further protection to the surface of the skull. One-minute pre-cooling and two-minute post cooling was applied before and after FUS exposures. Temporal temperature evaluation of the brain, skull, and the other tissue were calculated based on Pennes Bioheat Equation. Both acoustic pressure and temperature distribution on the 3D numerical model were finally acquired to evaluate the targeting accuracy and thermal safety.

### MRI imaging

All the NHPs were placed on an MRI compatible stereotaxic device for MRI imaging at 3T Siemens scanner, and an 8-channel surface receiver array coil was used to acquire both structural and functional MRI images. Under 0.8-1.1% light anesthesia, the resting state functional MRI scans were performed on NHPs (M, N, O, P, Q, T) using a T2* weighted EPI sequence (TR=2000 ms; TE=28.2 ms; FA=70°; 1.65mm isotropic resolution, FOV=106 × 106 × 53mm^3^, 64 × 64× 32 matrix voxels, 456 volumes per run). The baseline scans were performed on all six NHPs including 2-4 runs (∼15 min per run, shown in Fig. 1a) for each NHP, but without FUS sonication. For the groups of NHPs undergoing FUS neuromodulation (NHPs O and P) and FUS BBB opening (NHPs M and T), a continuous 4 runs of resting state fMRI acquisitions were acquired at approximately 45 min after 2 min FUS exposure described in the previous section. In the same session of the functional scans, the T1 weighted structural scans were also acquired (TR = 2580 ms; TE = 2.81 ms; FA = 9°, isotropic 0.5 mm resolution; FOV 128 ×128 × 60mm).

To evaluate the safety and efficiency of BBB opening, additional T1 weighted structural scans (same protocol with the regular T1, named post T1 in Fig. 1a) were acquired 30 min after IV administration of Gadolinium contrast agent (0.2 ml/kg). In normal condition, the Gadolinium contrast cannot cross the intact BBB due to its relatively large molecular size, however with FUS-mediated BBB opening, the contrast agent was utilized as an efficient means to visualize the site of BBB opening due to its increasing effects on BBB permeability (Karakatsani et al., 2017).

### Data processing and analysis

All the fMRI data were processed using a pipeline combining FSL (FSL 6.0.3) and Matlab (Matlab 2019b, The MathWorks, Inc., Natick, MA, United States). Firstly, preprocessing steps including motion correction, slice-timer correlation and B0 field map distortion correction were applied using FSL 6.0.3 (Jenkinson et al., 2012). Both temporal and spatial filtering were applied: a high pass temporal filter with 100 s cutoff was applied to remove the low-frequency noise; and a spatial filter with 3 mm FWHM Gaussian kernel was applied to smooth the fMRI data. In addition, global signal regression was applied on each dataset to remove the global artifacts due to motion and respiration (Murphy & Fox, 2017; Wong, et al., 2013). Finally, linear registration was achieved from functional to structural images, and then from structural images to the standard D99 template of NHP brain (Reveley et al., 2017), using the FAST tools of FSL.

In order to calculate the functional connectivity of the NHPs, we calculated seed-based correlation between the chosen ROI of 3.5 × 3.5 × 3.5 mm^3^ and the other nodes or areas within the relevant brain networks including default mode network (DMN), frontotemporal network (FTN), salience network (SN) and visual networks (VN). For quantification of functional connectivity in cortical regions, we selected ROI located at right Caudate; for quantification of different brain networks, we selected ROI located at dorsolateral prefrontal cortex (DMN)), medial prefrontal cortex (FTN), insular cortex (SN) and primary visual cortex (VN). The central coordinates of the ROI adopted in this study are listed in Table S1. The procedures above are performed for every run of resting state fMRI of NHPs including 18 baseline runs, 8 runs of FUS neuromodulation and 8 runs of FUS-mediated BBB opening. Using Fisher’ Z transformation, the transformed correlation coefficients between the seed ROI and the other brain area were calculated. The resulting correlations in terms of Fisher’ Z scores were then fed into Kruskal-Wallis test (McKight & Najab, 2010) to compare the BOLD activity between baseline, FUS neuromodulation and FUS-mediated BBB opening. The statistical threshold was set as 0.05, and when there is a statistical significant difference between the three groups, a post-hoc non parametric permutation test (Nichols & Holmes, 2002) with 5000 resamples were performed to compare the correlations between every two groups, with p < 0.05 representing a significance difference.

## Acknowledgements

The authors wish to acknowledge the ZI-ICM team for the experiential setup and anesthesia procedures during MRI imaging, and Dr. Ray F. Lee for MRI sequence setup and data collection. The work is supported by NIH R01 MH112142, NIH R01 EB009041, NIH R01 AG038961 and BBRF YI grant 31298.

## References

Bongioanni, A., Folloni, D., Verhagen, L., Sallet, J., Klein-Flügge, M.C. and Rushworth, M.F., 2021. Activation and disruption of a neural mechanism for novel choice in monkeys. Nature, 591(7849), pp.270–274.

Banaie Boroujeni K, Sigona MK, Treuting RL, Manuel TJ, Caskey CF, Womelsdorf T. Anterior cingulate cortex causally supports flexible learning under motivationally challenging and cognitively demanding conditions. PLoS Biol. 2022 Sep 6;20(9):e3001785. doi: 10.1371/journal.pbio.3001785. PMID: 36067198; PMCID: PMC9481162.

Blackmore, J., Shrivastava, S., Sallet, J., Butler, C. R., & Cleveland, R. O. (2019). Ultrasound neuromodulation: a review of results, mechanisms and safety. Ultrasound in medicine & biology, 45(7), 1509–1536.

Deffieux, T. and Konofagou, E.E., 2010. Numerical study of a simple transcranial focused ultrasound system applied to blood-brain barrier opening. IEEE transactions on ultrasonics, ferroelectrics, and frequency control, 57(12), pp.2637–2653.

Deffieux, T., Younan, Y., Wattiez, N., Tanter, M., Pouget, P., & Aubry, J.-F. (2013). Low-intensity focused ultrasound modulates monkey visuomotor behavior. Current Biology, 23(23), 2430–2433.

Dobryakova, E., & Smith, D. V. (2022). Reward enhances connectivity between the ventral striatum and the default mode network. Neuroimage, 258, 119398.

Downs, M. E., Teichert, T., Buch, A., Karakatsani, M. E., Sierra, C., Chen, S., … Ferrera, V. P. (2017). Toward a cognitive neural prosthesis using focused ultrasound. Frontiers in Neuroscience, 11, 607.

Duck, F. A. (2013). Physical properties of tissues: a comprehensive reference book: Academic press.

Folloni, D., Verhagen, L., Mars, R. B., Fouragnan, E., Constans, C., Aubry, J.-F., … Sallet, J. (2019). Manipulation of subcortical and deep cortical activity in the primate brain using transcranial focused ultrasound stimulation. Neuron, 101(6), 1109-1116. e1105.

Fouragnan, E. F., Chau, B. K. H., Folloni, D., Kolling, N., Verhagen, L., Klein-Flugge, M., … Rushworth, M. F. S. (2019). The macaque anterior cingulate cortex translates counterfactual choice value into actual behavioral change. Nat Neurosci, 22(5), 797–808. doi:10.1038/s41593-019-0375-6

Garin, C. M., Hori, Y., Everling, S., Whitlow, C. T., Calabro, F. J., Luna, B., … Dhenain, M. (2022). An evolutionary gap in primate default mode network organization. Cell reports, 39(2), 110669.

Goss, S.A., Frizzell, L.A. and Dunn, F., 1979. Ultrasonic absorption and attenuation in mammalian tissues. Ultrasound in medicine & biology, 5(2), pp.181–186.

Hasgall, P.A., Di Gennaro, F., Baumgartner, C., Neufeld, E., Gosselin, M.C., Payne, D., Klingenböck, A. and Kuster, N., (2022). IT’IS database for thermal and electromagnetic parameters of biological tissues, version 4.1. doi:10.13099/VIP21000-04-1

Jenkinson, M., Beckmann, C. F., Behrens, T. E., Woolrich, M. W., & Smith, S. M. (2012). Fsl. Neuroimage, 62(2), 782–790.

Karakatsani, M. E. M., Samiotaki, G. M., Downs, M. E., Ferrera, V. P., & Konofagou, E. E. (2017). Targeting Effects on the Volume of the Focused Ultrasound-Induced Blood-Brain Barrier Opening in Nonhuman Primates In Vivo. IEEE Trans Ultrason Ferroelectr Freq Control, 64(5), 798–810. doi:10.1109/TUFFC.2017.2681695

Kubanek, J. (2018). Neuromodulation with transcranial focused ultrasound. Neurosurg Focus, 44(2), E14. doi:10.3171/2017.11.FOCUS17621

Lv, H., Wang, Z., Tong, E., Williams, L.M., Zaharchuk, G., Zeineh, M., Goldstein-Piekarski, A.N., Ball, T.M., Liao, C. and Wintermark, M., 2018. Resting-state functional MRI: everything that nonexperts have always wanted to know. American Journal of Neuroradiology, 39(8), pp.1390–1399.

Mantini, D., Gerits, A., Nelissen, K., Durand, J.-B., Joly, O., Simone, L., … Buckner, R. L. (2011). Default mode of brain function in monkeys. Journal of Neuroscience, 31(36), 12954–12962.

McDannold, N., Arvanitis, C. D., Vykhodtseva, N., & Livingstone, M. S. (2012). Temporary Disruption of the Blood–Brain Barrier by Use of Ultrasound and Microbubbles: Safety and Efficacy Evaluation in Rhesus MacaquesBlood–Brain Barrier Disruption via Focused Ultrasound. Cancer research, 72(14), 3652–3663.

McKight, P. E., & Najab, J. (2010). Kruskal-wallis test. The corsini encyclopedia of psychology, 1–1.

Meng, Y., Pople, C. B., Lea-Banks, H., Abrahao, A., Davidson, B., Suppiah, S., … Hynynen, K. (2019a). Safety and efficacy of focused ultrasound induced blood-brain barrier opening, an integrative review of animal and human studies. Journal of Controlled Release, 309, 25–36.

Meng, Y., MacIntosh, B. J., Shirzadi, Z., Kiss, A., Bethune, A., Heyn, C., … Lipsman, N. (2019b). Resting state functional connectivity changes after MR-guided focused ultrasound mediated blood-brain barrier opening in patients with Alzheimer’s disease. Neuroimage, 200, 275–280. doi:10.1016/j.neuroimage.2019.06.060

Moyano, D. B., Paraiso, D. A., & González-Lezcano, R. A. (2022). Possible Effects on Health of Ultrasound Exposure, Risk Factors in the Work Environment and Occupational Safety Review. Paper presented at the Healthcare.

Munoz, F., Meaney, A., Gross, A., Liu, K., Pouliopoulos, A., Liu, D., … Ferrera, V. (2022). Long term study of motivational and cognitive effects of low-intensity focused ultrasound neuromodulation in the dorsal striatum of nonhuman primates. Brain Stimulation, 15(2), 360–372.

Murphy, K., & Fox, M. D. (2017). Towards a consensus regarding global signal regression for resting state functional connectivity MRI. Neuroimage, 154, 169–173.

Nichols, T. E., & Holmes, A. P. (2002). Nonparametric permutation tests for functional neuroimaging: a primer with examples. Human brain mapping, 15(1), 1–25.

Nitsche, M. A., Cohen, L. G., Wassermann, E. M., Priori, A., Lang, N., Antal, A., … Fregni, F. (2008). Transcranial direct current stimulation: state of the art 2008. Brain Stimulation, 1(3), 206–223.

Pouliopoulos, A. N., Kwon, N., Jensen, G., Meaney, A., Niimi, Y., Burgess, M. T., … Kamimura, H. A. (2021). Safety evaluation of a clinical focused ultrasound system for neuronavigation guided blood-brain barrier opening in non-human primates. Scientific reports, 11(1), 1–17.

Rao, B., Xu, D., Zhao, C., Wang, S., Li, X., Sun, W., … Xu, H. (2021). Development of functional connectivity within and among the resting-state networks in anesthetized rhesus monkeys. Neuroimage, 242, 118473.

Reveley, C., Gruslys, A., Ye, F. Q., Glen, D., Samaha, J., b, E. R., … Saleem, K. S. (2017). Three-Dimensional Digital Template Atlas of the Macaque Brain. Cereb Cortex, 27(9), 4463–4477. doi:10.1093/cercor/bhw248

Sherman, S. M. (2016). Thalamus plays a central role in ongoing cortical functioning. Nature neuroscience, 19(4), 533–541.

Ter Haar, G. and Coussios, C., 2007. High intensity focused ultrasound: past, present and future. International Journal of Hyperthermia, 23(2), pp.85–87.

Todd, N., Zhang, Y., Arcaro, M., Becerra, L., Borsook, D., Livingstone, M., & McDannold, N. (2018). Focused ultrasound induced opening of the blood-brain barrier disrupts inter-hemispheric resting state functional connectivity in the rat brain. Neuroimage, 178, 414–422. doi:10.1016/j.neuroimage.2018.05.063

Todd, N., Zhang, Y., Livingstone, M., Borsook, D., & McDannold, N. (2019). The neurovascular response is attenuated by focused ultrasound-mediated disruption of the blood-brain barrier. Neuroimage, 201, 116010. doi:10.1016/j.neuroimage.2019.116010

Treeby, B.E. and Cox, B.T., 2010. k-Wave: MATLAB toolbox for the simulation and reconstruction of photoacoustic wave fields. Journal of biomedical optics, 15(2), pp.021314–021314.

Tufail, Y., Matyushov, A., Baldwin, N., Tauchmann, M. L., Georges, J., Yoshihiro, A., … Tyler, W. J. (2010). Transcranial pulsed ultrasound stimulates intact brain circuits. Neuron, 66(5), 681–694.

Verhagen, L., Gallea, C., Folloni, D., Constans, C., Jensen, D. E., Ahnine, H., … Sallet, J. (2019). Offline impact of transcranial focused ultrasound on cortical activation in primates. Elife, 8. doi:10.7554/eLife.40541

Walsh, V., & Cowey, A. (2000). Transcranial magnetic stimulation and cognitive neuroscience. Nature Reviews Neuroscience, 1(1), 73–80.

Wong, C. W., Olafsson, V., Tal, O., & Liu, T. T. (2013). The amplitude of the resting-state fMRI global signal is related to EEG vigilance measures. Neuroimage, 83, 983–990.

Wu, S. Y., Aurup, C., Sanchez, C. S., Grondin, J., Zheng, W., Kamimura, H., … Konofagou, E. E. (2018). Efficient Blood-Brain Barrier Opening in Primates with Neuronavigation-Guided Ultrasound and Real-Time Acoustic Mapping. Sci Rep, 8(1), 7978. doi:10.1038/s41598-018-25904-9

Xu, T., Sturgeon, D., Ramirez, J. S., Froudist-Walsh, S., Margulies, D. S., Schroeder, C. E., … Milham, M. P. (2019). Interindividual variability of functional connectivity in awake and anesthetized rhesus macaque monkeys. Biological Psychiatry: Cognitive Neuroscience and Neuroimaging, 4(6), 543–553.

Zhang, Z., Cai, D. C., Wang, Z., Zeljic, K., Wang, Z., & Wang, Y. (2019). Isoflurane-induced burst suppression increases intrinsic functional connectivity of the monkey brain. Frontiers in neuroscience, 13, 296.

