## Supplemental Materials for "Alteration of functional connectivity in the cortex and major brain networks of non-human primates following focused ultrasound exposure"

**Table S1** Seed ROI selected for measurement of functional connectivity, with all the coordinates defined in D99 atlas (V 1.2b, 0.25mm isotropic resolution)

| Area | Region/Description | Centroid Voxel Location | | | Centroid coordinate[1] (mm) | | |
|---|---|---|---|---|---|---|---|
| | | X | Y | Z | X | Y | Z |
| 8Ad, right brain | Dorsal lateral Prefrontal Cortex | 81 | 238 | 161 | -14 | 9.75 | 12.75 |
| 46d, right brain | Dorsal lateral Prefrontal Cortex | 92 | 272 | 148 | -11.25 | 18.25 | 9.5 |
| 8Bs, right brain | Dorsal lateral Prefrontal Cortex | 95 | 236 | 158 | -10.5 | 9.25 | 12 |
| 9m | Medial Prefrontal Cortex | 137 | 289 | 147 | 0 | 22.5 | 9.25 |
| 23b | Posterior Cingulate Cortex | 137 | 143 | 165 | 0 | -14 | 13.75 |
| Ial, right brain | Insular Cortex | 82 | 219 | 89 | -13.75 | 5 | -5.25 |
| V1, left brain | Primary Visual Cortex | 185 | 51 | 143 | 12 | -37 | 8.25 |

[1] Coordinates (0,0,0) represents the center of the D99 brain atlas.

**Table S2** Acoustic and bio-thermal parameters in k-wave simulation

| Tissue Properties | Symbol | Skull | Brain | Muscle | Water |
|---|---|---|---|---|---|
| Speed of sound[1] (m/s) | $c$ | $c_0 + (c_{max} - c_0)\rho_{CT}$ | 1546.3 | 1588.4 | 1482.3 |
| Density[2] ($kg/m^3$) | $\rho$ | $\rho_0 + (\rho_{max} - \rho_0)\rho_{CT}$ | 1046 | 1090 | 998.2 |
| Specific heat (J/(kg·K)) | $C$ | 1313 | 3630 | 3421 | 4178 |
| Thermal conductivity (W/(m·K)) | $k$ | 0.3 | 0.51 | 0.49 | 0.6 |
| Acoustic attenuation[3] (db/cm) | $\alpha$ | $\alpha_0 + (\alpha_{max} - \alpha_0)f^{1.1}\Psi^{\beta}$ | 0.59*$f^{1.1}$ | 0.62*$f^{1.1}$ | 0.0022$*$ $f^{1.1}$ |
| Perfusion rate ($s^{-1}$) | $\omega$ | 0 | 0.0098 | 0.0006 | 0 |

[1] $c_0 = 1854.7\ m/s,\ \ c_{max} = 3360\ m/s.\ \rho_{CT} = \frac{HU}{\max(HU)}$ is normalized CT Hounsfield Unit

[2] $\rho_0 = 1080\ kg/m^3;\ \ \rho_{max} = 2100\ kg/m^3.$

[3] $f = 0.5\ MHz$, $\alpha_0 = 0.2\ db/(cm \cdot MHz^{-1.1})$, $\alpha_{max} = 8\ db/(cm \cdot MHz^{-1.1})$, $\Psi = 1 - \rho_{CT}$ is the porosity and $\beta = 0.5$. The absorption coefficient of the skull is set as $\alpha_{abs} = 2.7\ (db/(cm \cdot MHz^{-1.1})$ during thermal simulation.

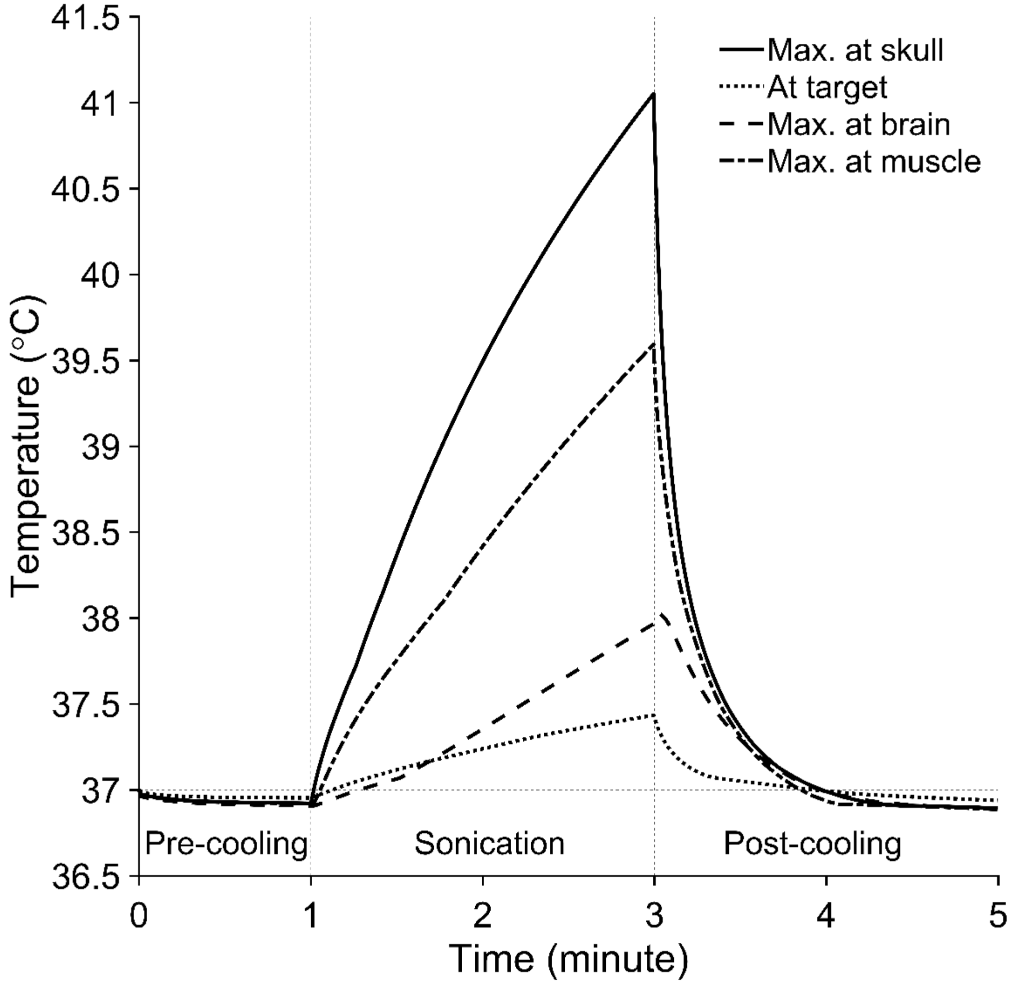


**Fig.S1** Temporal temperature curves of the target, maximum temperature in the skull, brain and muscle. The thermal simulation is running in k-wave, with the following ultrasound parameters: source PNP=115 kPa, target PNP=800 kPa, 2 Hz PRF, 2% duty cycle and 10 ms pulse width and 2 min total sonication. One-minute pre-cooling and two-minute post-cooling were applied to protect the skull from thermal damage.

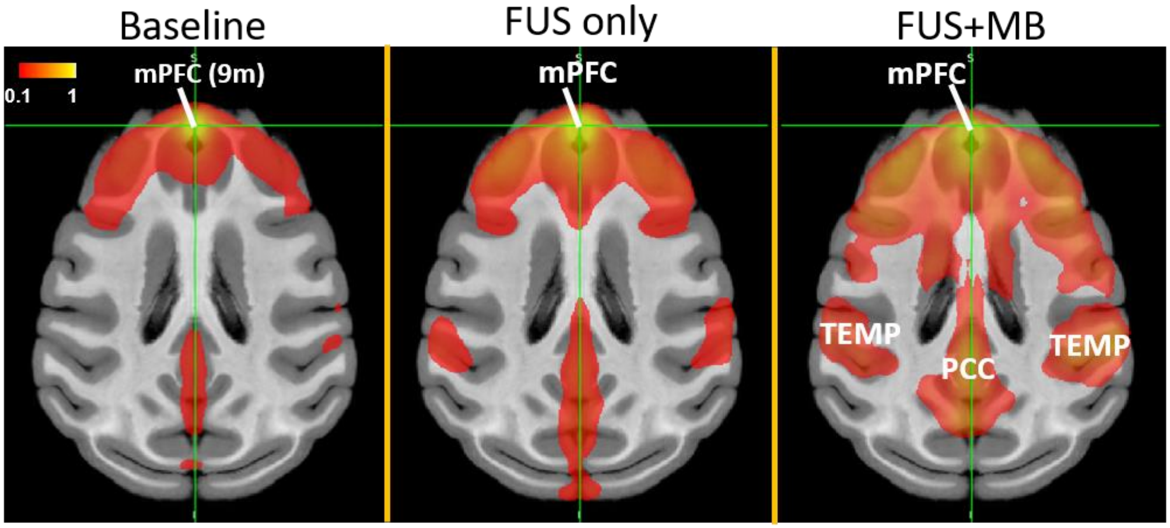


**Fig.S2** Seed-based connectivity maps of resting state fMRI in the horizontal plane without FUS (baseline), after FUS (FUS only) and FUS with microbubbles (FUS+MB). The red-yellow color maps indicate Fisher z-scores with seeds in the area 9m of mPFC. When applying FUS with microbubbles, strong activation was found between mPFC and TEMP, and mPFC and PCC. mPFC: medial prefrontal cortex, TEMP: temporal lobe, PCC: posterior cingulate cortex.

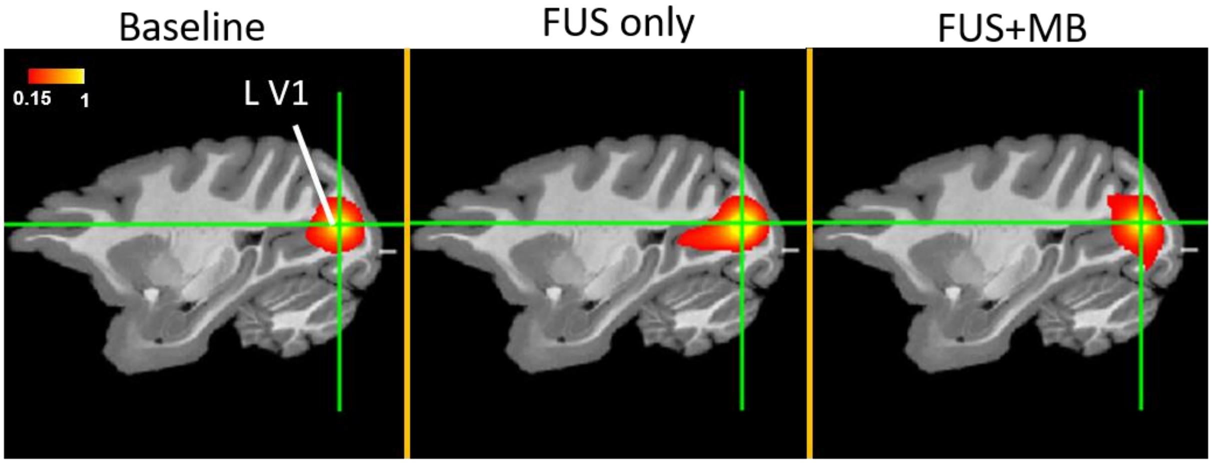


**Fig.S3** Seed-based correlation maps of resting state fMRI without FUS (baseline), after FUS exposure (FUS only) and after FUS with microbubbles (FUS+MB). The red-yellow color maps indicate Fisher z-scores with the seed ROI in the left primary visual cortex (LV1). Compared to baseline, both FUS and FUS with microbubbles did not show a significant alteration of the functional connectivity between left primary visual cortex and the other brain regions.

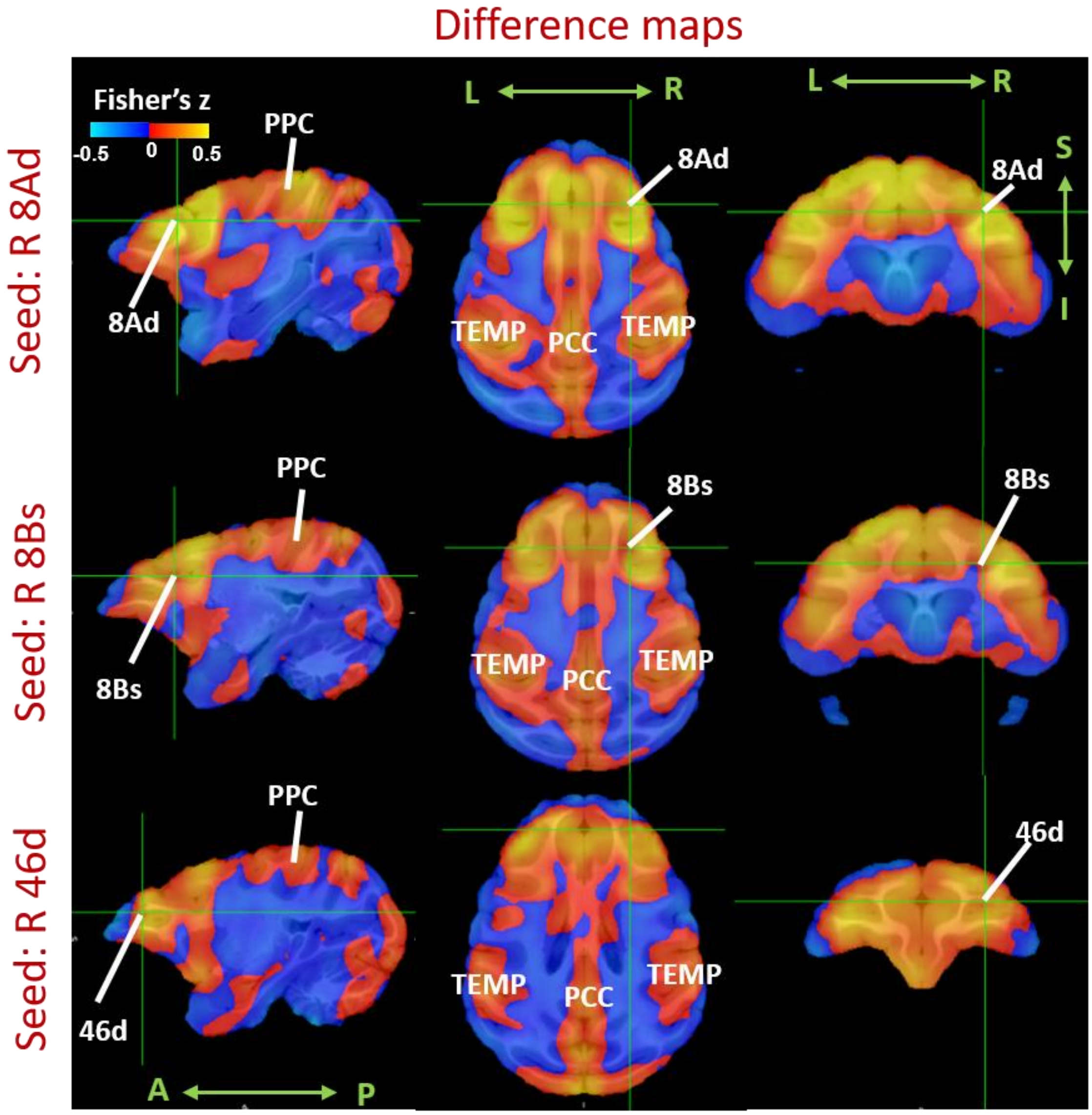


**Fig.S4** Seed-Based correlation difference maps between FUS exposure and Baselines, with the seed chosen as right 8Ad, right 8Bs and right 46d respectively. The dlPFC-PCC and dlPFC-TEMP were activated by FUS exposure, no matter which seed in dlPFC was chosen. dlPFC: dorsolateral prefrontal cortex, PCC: posterior cingulate cortex. TEMP: temporal lobe.
